# Intranasal Anti-CD3 Antibody Treatment Attenuates Post–COVID Neuroinflammation and Enhances Hippocampal Neurogenesis and Cognitive Function in Mice

**DOI:** 10.64898/2026.04.07.716934

**Authors:** Peiwen Lu, Saef Izzy, Patrick Da Silva, Harm Tjebbe Imkamp, Jonathan R. Christenson, Taha Yahya, Maryam Hazim Al Mansi, Amir Alawi, Thais G. Moreira, Michelle Monje, Howard L. Weiner, Akiko Iwasaki

## Abstract

Cognitive impairment is a disabling feature of Long COVID, with data supporting neuroinflammation and maladaptive glial responses as primary drivers. Nasal administration of an anti-CD3 monoclonal antibody (aCD3 mAb) has shown therapeutic benefits in autoimmune and CNS disease models. Using a respiratory-restricted mild SARS-CoV-2 mouse model of Long COVID, we show that nasal anti-CD3 mAb, administered shortly after infection or during chronic neuroinflammation, increased brain FoxP3+ IL–10+ Tregs, reduced microglial and astrocytic gliosis in the white matter and hippocampus, restored neurogenesis, and improved short-term memory. Nasal aCD3 mAb reprogrammed microglia from an antigen-presenting, NF-κB-driven inflammatory state toward chemokine signaling, phagosome, and TGF β–related regulatory phenotype. Patients with Long COVID with neurological symptoms had lower circulating Treg populations. These findings identify nasal administration of aCD3 mAb as a noninvasive strategy to control neuroinflammation, restore the neurogenic niche, and offer a novel approach to treating cognitive impairment in Long COVID.

## Introduction

Long COVID, or post-acute sequelae of SARS CoV-2 infection, is a heterogeneous syndrome in which symptoms persist or recur months after the initial illness, typically beyond the period of viral clearance (Groff et al., 2021; Nalbandian et al., 2021). Cognitive and neuropsychiatric manifestations, including impaired attention, memory, and executive function that are commonly described as ‘brain fog,’ together with fatigue and sleep disturbances, are among its most debilitating features (Monje and Iwasaki, 2022; Xu et al., 2022). These symptoms can persist for years, substantially limiting work capacity, daily functioning, and social participation, often in adults during mid-life (Al-Aly et al., 2024; Davis et al., 2023). Understanding how post-viral CNS dysfunction contributes is crucial, not only for persistent disability but also for processes associated with brain aging, neurodegeneration, and increased vulnerability to cognitive decline in later life decline in at-risk individuals(Bonhenry et al., 2024; Bruno et al., 2025; Seo et al., 2025).

However, the biological mechanisms linking respiratory viral infection to sustained cognitive impairment remain poorly defined, and no approved therapies target the neurological manifestations of Long COVID (Iwasaki and Putrino, 2023; Peluso and Deeks, 2024). Progress has been hindered by two major challenges: the heterogeneous pathophysiology of Long COVID and the lack of preclinical models that faithfully recapitulate persistent CNS dysfunction after respiratory infection (Ellen, 2025; Hamlin and Blish, 2024; Jansen et al., 2022; Pagliano et al., 2025), leaving clinical trials to proceed without clear biomarkers, patient stratification, or strong mechanistic support. Available interventions also remain limited once symptoms are established. Although vaccination attenuates acute disease severity and reduces the risk of developing Long COVID by lowering infection (Green et al., 2025; Vanderheiden et al., 2024), protection is incomplete, breakthrough infections still occur, and vaccination does not consistently alter the severity or profile of neurological symptoms in individuals with established symptoms(Mukherjee et al., 2025; Notarte et al., 2022). These limitations underscore the need for mechanism-based therapies that directly target neuroimmune processes sustaining chronic CNS dysfunction.

Multiple studies indicate that SARS-CoV-2 infection can lead to sustained not only peripheral immune dysregulation(Yin et al., 2024) but also neuroinflammation, microglial reactivity, blood-brain barrier disruption, and impaired hippocampal neurogenesis in the CNS, even when acute infection is restricted to the respiratory tract(Braga et al., 2023; Fujimoto et al., 2025; Greene et al., 2024; Saikarthik et al., 2025). Experimental models of mild, respiratory-restricted SARS-CoV-2 infection recapitulates these features, revealing persistent microglial reactivity, white matter pathology, oligodendrocyte and myelin loss, as well as chemokine changes including elevated CCL11 that impair hippocampal neurogenesis and mirror alterations observed in people with Long COVID(Fernandez-Castaneda et al., 2022; Israelow et al., 2020).

Foxp3+ regulatory T (Treg) cells are a subset of CD4+ T cells that maintain immune homeostasis by restraining excessive effector responses and producing anti-inflammatory mediators such as IL-10 and TGF-β (Vignali et al., 2008). Across diverse neuroinflammatory and neurodegenerative conditions, Foxp3+ Treg cells have emerged as key regulators that limit CNS inflammation and support tissue repair, and their quantitative or functional impairment has been linked to worse neurological outcomes.(Liston et al., 2022; Liston et al., 2024). These observations suggest that augmenting Treg–mediated regulation could counteract the neuroinflammatory component of Long COVID and promote recovery of CNS function.

Mucosal delivery provides a physiologic, tolerogenic route for inducing regulatory T cell responses and has shown a favorable safety profile in prior studies(Ochi et al., 2006; Weiner et al., 2011). Nasal administration of anti-CD3 monoclonal antibody (aCD3 mAb) induces Treg responses(Kuhn and Weiner, 2016; Wu et al., 2008) and has shown therapeutic benefit in both autoimmune and CNS models of disease, including models of brain injury, progressive multiple sclerosis (MS) and Alzheimer’s disease (Izzy et al., 2025; Lopes et al., 2023; Mayo et al., 2016). Nasal human aCD3 mAb is currently being investigated in clinical trials across several high-need therapeutic areas, including MS and Alzheimer’s disease(Chitnis et al., 2026; Singhal et al., 2025). We therefore asked whether nasal aCD3 could influence outcomes in model of Long COVID by modulating CNS immune responses.

Here, we used a validated mild COVID mouse model in which SARS-CoV-2 infection is restricted to the respiratory tract yet leads to persistent CNS pathology that resembles the human Long COVID phenotype. We show that nasal aCD3 induces Foxp3+ Treg cells that access the CNS, reprogram microglia, reduce gliosis across affected regions, and restore hippocampal neurogenesis and cognitive performance, thereby directly ameliorating Long COVID–like pathology. Together, these findings identify nasal aCD3–mediated Treg induction as a mechanistically grounded, immune-based strategy for treating Long COVID–associated neurological manifestations and, more broadly, for targeting neuroinflammation in post-infectious brain disorders.

## Results

### Early nasal anti-CD3 mAb treatment limits neuroinflammatory changes in a Long COVID mouse model

To test whether aCD3 mAb can modify post-infectious CNS inflammation, we administered aCD3 mAb intranasally in a validated mild COVID model of respiratory-restricted SARS-CoV-2 infection (Figure 1A)(Fernandez-Castaneda et al., 2022). Although aCD3 engages the T cell receptor complex (Chatenoud and Bluestone, 2007), 4 weeks of treatment did not alter anti–SARS CoV-2 spike antibody titers, indicating that nasal anti-CD3 did not blunt the antiviral humoral immunity (Figure S1A). We next quantified glial responses in brain regions implicated in post-infectious cognitive dysfunction, including subcortical white matter and hippocampus. Infected mice receiving isotype control showed a marked increase in IBA1+ microglia in both regions compared with mock-infected animals, consistent with the previous report (Fernandez-Castaneda et al., 2022)(Figure 1B). Nasal aCD3 substantially reduced microglial accumulation in each region, restoring cell numbers toward mock levels (Figure 1B, S1B). We observed a similar pattern for astrocytic gliosis. GFAP immunostaining revealed robust astrocyte reactivity in the subcortical white matter and hippocampus of isotype-treated mild COVID mice (Eng et al., 1971; Hol and Pekny, 2015), whereas aCD3 treatment significantly reduced GFAP+ astrocyte density across these regions (Figure 1C). To determine whether these CNS changes were accompanied by systemic immune modulation, we profiled circulating cytokines and chemokines. Nasal aCD3 decreased serum CXCL10 and increased IL-10 relative to isotype treated mild COVID mice, consistent with a shift toward a more regulatory immune milieu (Figure. 1D). Together, these data show that initiating nasal aCD3 soon after acute SARS CoV-2 infection attenuates microglial and astrocytic activation, prevents the establishment of chronic neuroinflammation and is associated with systemic cytokine changes indicative of enhanced immune regulation.

**Figure. 1.**
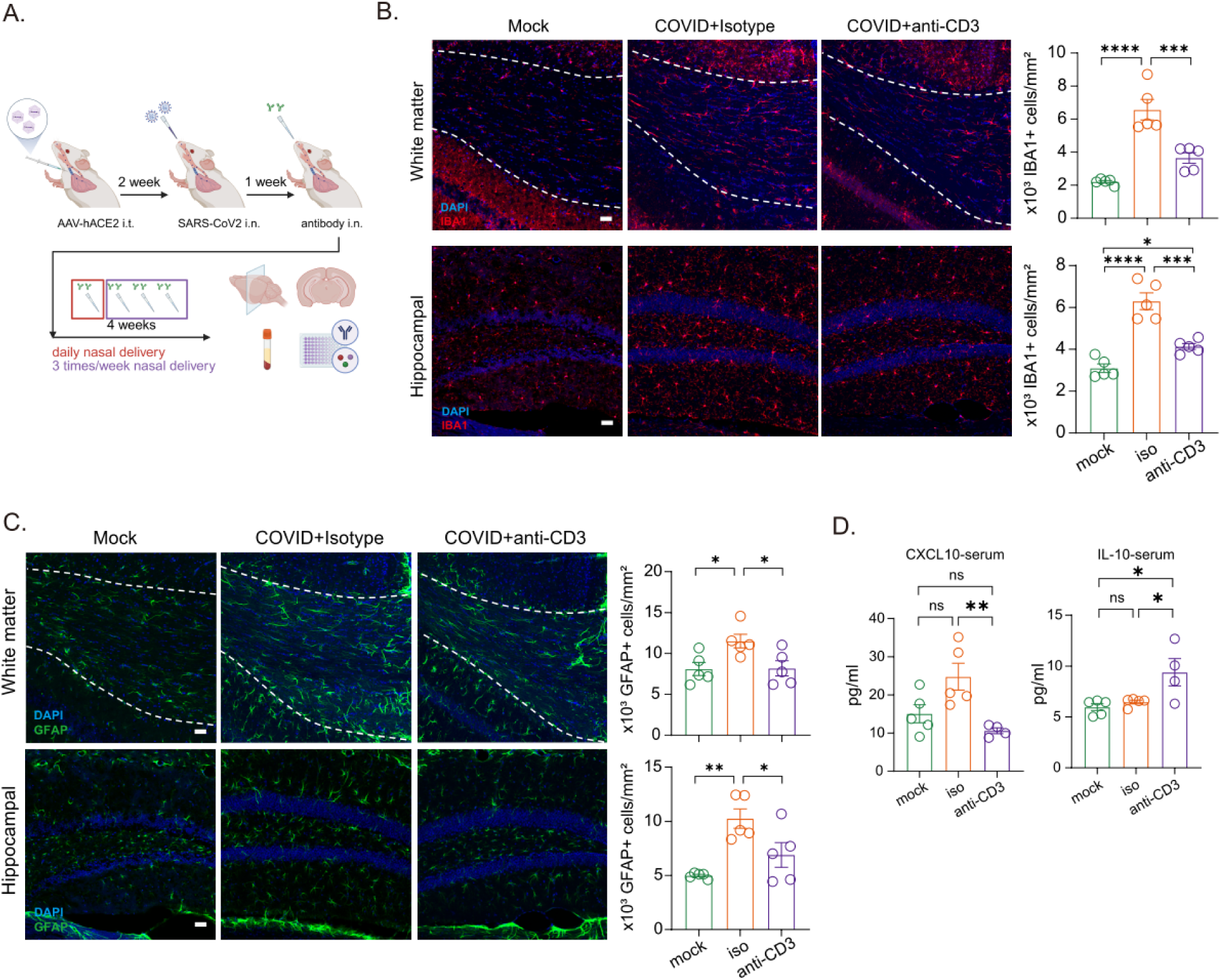
Nasal anti-CD3 treatment reduces CNS neuroinflammation after acute SARS CoV-2 infection. **(A**) Experimental paradigm for intranasal anti-CD3 administration in the mild COVID mouse model. Mice expressing human ACE2 received respiratory restricted SARS CoV-2 infection, followed by nasal delivery of isotype control or anti-CD3 antibody for 4 weeks. **(B)** Representative confocal images and quantification of IBA1+ microglia in subcortical white matter and dentate gyrus from mock mice, mild COVID mice treated with isotype control or anti-CD3; n=5 mice per group. **(C)** Representative confocal images and quantification of GFAP+ astrocytes in the same regions and groups; n = 5 mice per group. Scale bars, 100μm. **(D)** Serum level of CXCL10 and IL-10 in mock, mild COVID + isotype and mild COVID + anti-CD3 groups; n=4-5 mice per group. Data are mean ± SEM; each dot represents an individual mouse. \**p* <0.05, ** *p* <0.01, \*\*\**p* <0.001, \*\*\*\**p* <0.0001 (one way ANOVA followed by Tukey’s multiple comparison test).

### Early nasal anti-CD3 mAb reprograms microglial transcriptional states in the CNS

To define how nasal aCD3 mAb reshapes microglial states after mild COVID, we performed bulk RNA seq on subcortical microglia isolated from mock-infected mice and from mild COVID mice treated with isotype control or nasal aCD3 mAb (Figure 2A, S1C, S1D). Compared with mock, isotype-treated mild COVID mice showed extensive transcriptional reprogramming, with 209 differentially expressed genes (DEGs), whereas aCD3 treatment yielded 242 DEGs relative to isotype, indicating a robust shift in the infection-primed microglial transcriptome (Figure 2B). Analysis of genes that were significantly changed in both groups (mock versus isotype and aCD3 versus isotype) revealed a shared gene set that was strongly upregulated in microglia from isotype-treated mild COVID mice but returned toward mock levels in anti–CD3–treated mice, indicating that nasal aCD3 reverses an infection induced microglial activation program (Figure S1E)(Grant et al., 2024). Analysis of the full microglial DEG set highlighted multiple biological programs altered by infection and aCD3 treatment. This broader view included changes in immune-associated features, including antigen processing and presentation genes (e.g., Cd74, H2-Ab1, H2-K2) and chemokine/chemokine receptor-related genes (e.g., Ccl12, Ccr2) (Figure 2C). Gene ontology analysis of DEGs shared across groups revealed enrichment for pathways related to protein catabolism, autophagosome assembly and macrophage differentiation (Figure 2D). These signatures suggested that post-infection aCD3 treatment influences not only inflammatory signaling but also microglial phagocytic and structural interactions with myelinated axons. When we focused specifically on genes upregulated by aCD3 relative to isotype, we observed enrichment of pathways involved in chemokine production and chemokine receptor signaling, type I interferon and antiviral responses, antigen processing and presentation, and phagocytosis/endocytosis (Figure 2E). Thus, in this context, “microglial remodeling” reflects a coordinated re-balancing of microglial programs: nasal aCD3 enhances transcriptional pathways that support immune regulation, debris clearance, and maintenance of homeostasis in mice post-acute infection. However, this early intervention window is unlikely to be practical for clinical implementation for many individuals with Long COVID, who typically present months or even years after initial infection (Bonilla et al., 2023).

**Figure. 2.**
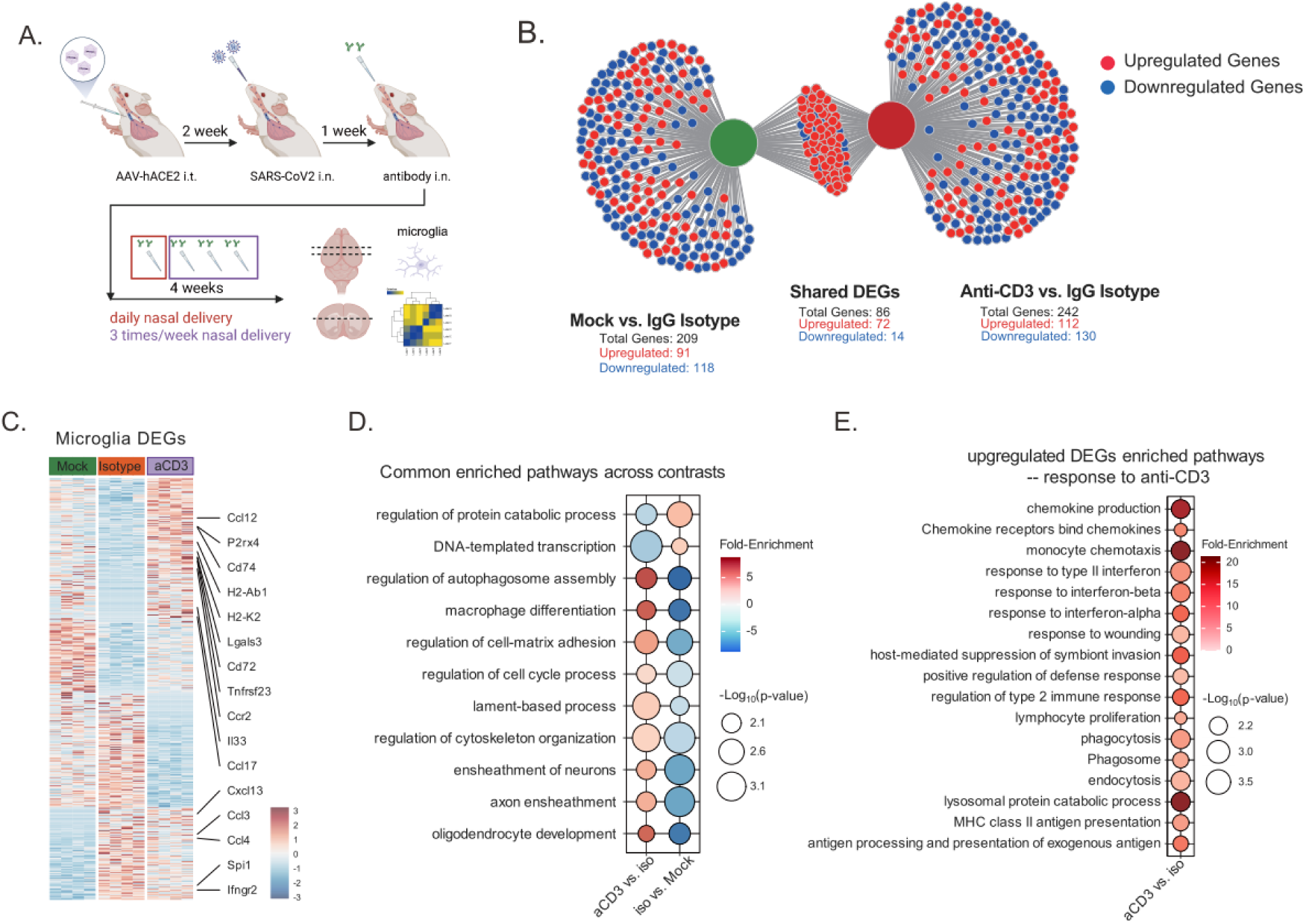
Nasal anti-CD3 treatment reprograms microglial transcriptional states in mild COVID. **(A**) Experimental paradigm for bulk RNA-seq of subcortical microglia from mock infected mice and mild COVID mice treated with isotype control or nasal anti-CD3. **(B)** Network plot showing overlap of differentially expressed genes (DEGs) in microglia from isotype treated mild COVID versus mock and anti CD3–treated mild COVID versus isotype; shared DEGs represent genes differentially expressed in both contrasts (associated Fig S1D). **(C)** Heatmap of microglial DEGs identified in either comparison (mock vs isotype or anti-CD vs isotype), n=4-5 mice per group. Selected genes associated with antigen presentation and chemokine signaling are highlighted. **(D)** Gene ontology (GO) categories enriched among cross comparison DEGs, illustrating pathways most consistently modulated by infection and treatment. **(E)** GO pathways enriched among genes upregulated in anti-CD3 treated versus isotype treated mild COVID microglia.

### Delayed nasal aCD3 mAb induces brain Treg cells and reverses established neuroinflammation in chronic post COVID setting

To test whether nasal aCD3 can treat, rather than prevent, post-infectious CNS pathology, we infected mice with SARS-CoV-2, allowed neuroinflammation to evolve for 4 weeks, and then initiated nasal aCD3 or isotype control treatment (Figure 3A). Delayed nasal aCD3 administration did not compromise antiviral immunity, as serum anti-spike IgG titers were comparable between aCD3– and isotype-treated infected mice (Figure 3B). We next assessed whether late aCD3 could still modify established gliosis. Quantitative immunostaining showed that IBA1+ microglia and GFAP+ astrocytes remained elevated in subcortical white matter and hippocampus of isotype-treated post-COVID mice, whereas aCD3 markedly reduced microglial and astrocytic cell numbers across these regions, approaching mock levels (Figure 3C, 3D, S2A). Thus, nasal aCD3 mAb was able to attenuate chronic gliosis even when initiated weeks after infection.

**Figure. 3.**
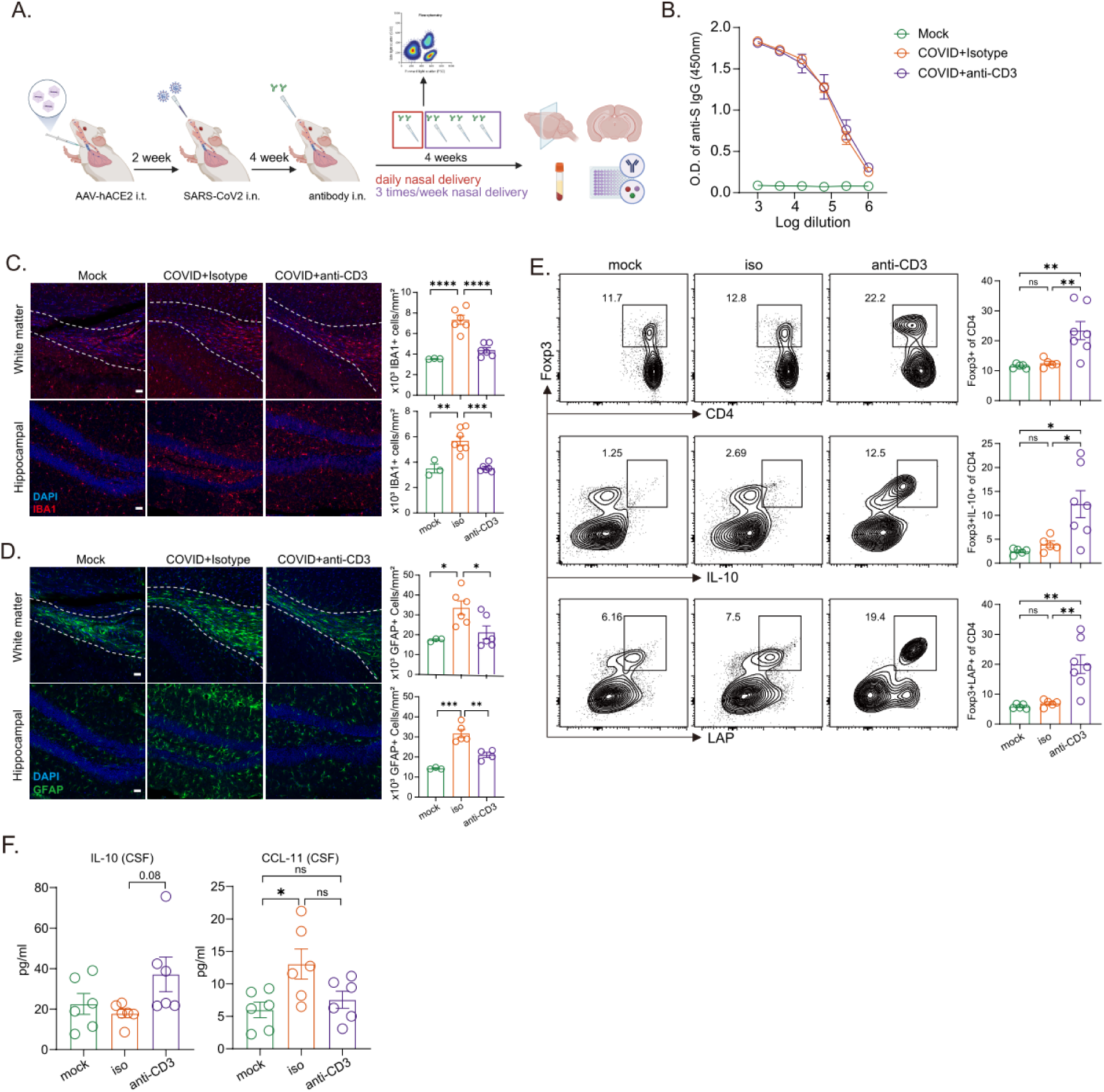
Nasal anti-CD3 induces brain Treg cells and reverses persistent neuroinflammation in mild COVID. **(A**) Experimental paradigm for delayed intranasal anti-CD3 treatment in the mild COVID mouse model. **(B)** Serum anti-spike IgG titers measured by ELISA, n=6-7 mice per group. (**C)** Representative confocal images and quantification of IBA1+ microglia in subcortical white matter and dentate gyrus from mock, mild COVID + isotype and mild COVID + aCD3 mice; n = 3–7 mice per group. **(D)** Representative confocal images and quantification of GFAP+ astrocytes in the same regions and groups; n=3-7 mice per group. Scale bars, 100μm. **(E)** Representative flow cytometry plots and quantification of Foxp3+, Foxp3+IL-10+, and Foxp3+LAP+ CD4 T cells (gated on live CD45+CD11b-TCRβ+CD4+) in brain tissue. **(F)** Cerebrospinal fluid (CSF) concentrations of IL-10 and CCL-11 in mock, mild COVID + isotype or anti-CD3 groups; n=6-7 mice per group. Data are mean ± SEM; each dot represents an individual mouse. \**p* <0.05, ** *p* <0.01, \*\*\**p* <0.001, \*\*\*\**p* <0.0001 (one way ANOVA followed by Tukey’s multiple comparison test).

Because Treg cells can restrain neuroinflammation through IL 10–dependent interactions with glia (Ito et al., 2019; Liston et al., 2022), we asked whether therapeutic nasal aCD3 expands brain Treg populations in post COVID setting as in other neuroinflammation mouse models (Izzy et al., 2025; Mayo et al., 2016). Flow cytometric analysis of brain CD4+ T cells after one week of treatment revealed that aCD3 significantly increased the frequency of Foxp3+ CD4+ T cells compared with isotype-treated mild COVID mice (Figure 3E). Within this expanded Treg compartment, we observed higher proportions of IL-10–producing Foxp3+ cells and latency associated peptide (LAP)-expressing Foxp3+ cells, consistent with enrichment of functionally suppressive Treg subsets (Figure 3E)(Gandhi et al., 2010; Rubtsov et al., 2008).

We previously linked elevated CCL11 to neuroinflammation in the mild COVID model and in Long COVID patients with cognitive symptoms (Fernandez-Castaneda et al., 2022). Analysis of cerebrospinal fluid showed that delayed nasal aCD3 shifted this cytokine–chemokine milieu toward a more regulatory profile: IL-10 levels trended upward, while CCL11 concentrations were significantly reduced relative to isotype-treated post-COVID mice (Figure 3F). Together, these findings demonstrate that delayed nasal aCD3 remains effective when delivered after the acute pulmonary phase, inducing IL-10–expressing brain Treg cells, normalizing CSF CCL11, and dampening established microglial and astrocytic activation in mild COVID mice.

### Nasal anti-CD3 antibody treatment improves neurogenesis and cognitive function post-COVID

Because cognitive dysfunction is a common symptom of Long COVID (Hampshire et al., 2021), we next asked whether nasal aCD3-mediated immune rebalancing improves downstream neural outcomes by restoring hippocampal neurogenesis and memory performance. In line with prior work (Fernandez-Castaneda et al., 2022), isotype-treated post-COVID mice showed a marked reduction in Doublecortin (DCX)+ neuroblasts in the dentate gyrus compared with mock controls, indicating impaired ongoing hippocampal neurogenesis. Nasal aCD3 significantly increased DCX+ cell density in post-COVID mice, and this rescue was evident whether mAb was initiated early (1 week after infection) or at a delayed time point (4 weeks after infection), indicating that the treatment promotes hippocampal neurogenesis across a broad temporal window (Figure 4A). To link these cellular changes to brain-wide cell states and behavior, we focused subsequent experiments on the delayed treatment paradigm used for single-cell RNAseq and cognitive testing (Figure 4B). For scRNA-seq, cerebrum tissue containing cortex, corpus callosum, and hippocampus were dissociated from each group. We profiled 22,941 cells from isotype-treated mild COVID mice, 24,395 cells from aCD3-treated mild COVID mice, and 13,458 cells from mock controls and annotated 18 major cell populations based on canonical markers (Figure 4C, S2B). At the level of broad classes, glial population frequencies were similar across conditions (Figure S2C), whereas non-glial populations, particularly neuronal lineages, showed the largest shifts with infection and treatment (Figure S2D). Stratifying by neuronal subtype revealed that nasal aCD3 was associated with an expansion of immature neuron populations and a relative reduction of mature excitatory neurons compared with isotype-treated post-COVID mice, consistent with enhanced neuroblast production (Figure 4D).

**Figure. 4.**
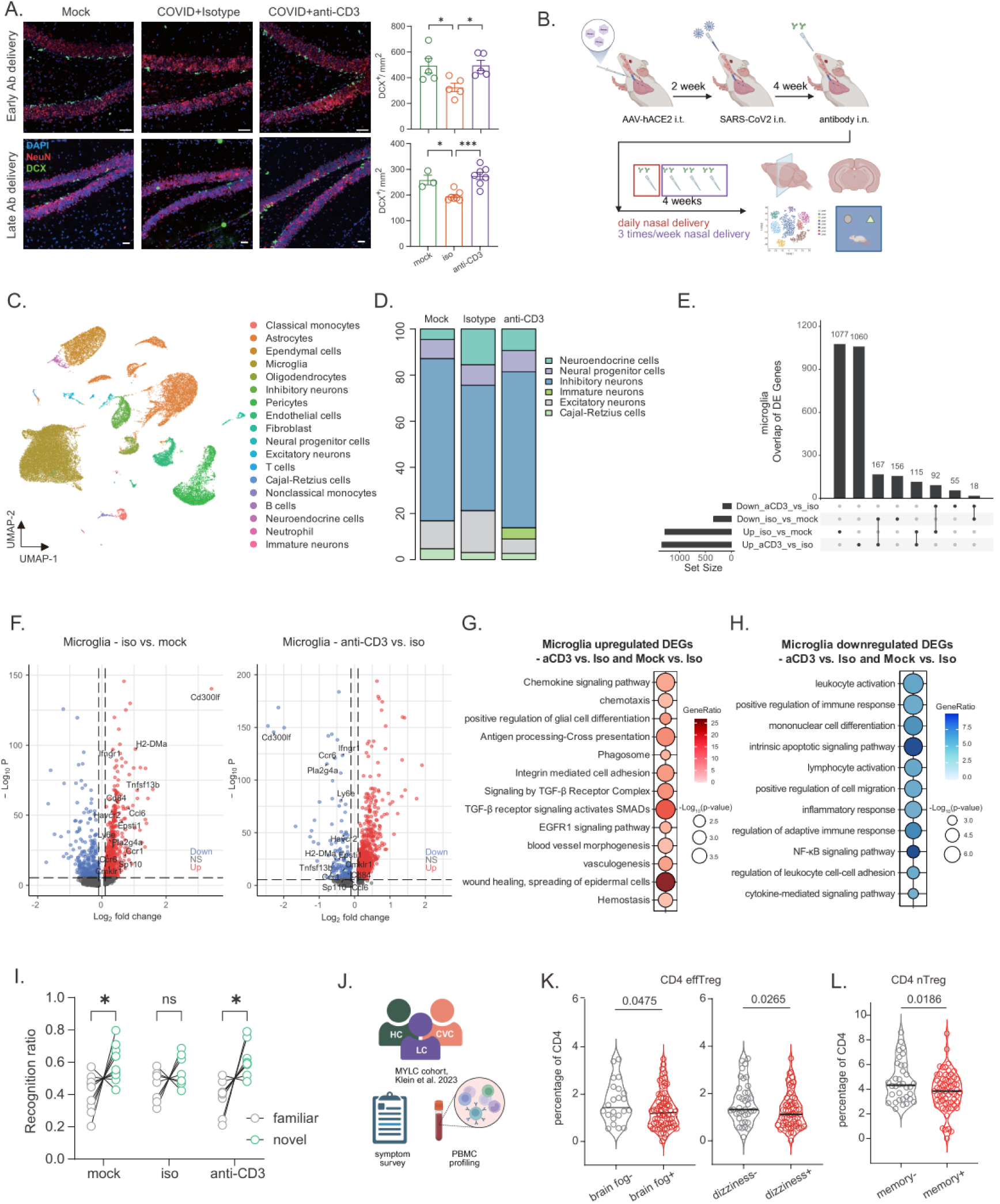
Nasal anti-CD3 antibody treatment improves neurogenesis and associated cognitive function in mild COVID mouse model. **(A)** Quantification and confocal micrograph of neuroblast (DCX+) in the dentate after 4 weeks anti-CD3 antibody administration in mild COVID or mock mice. Scale bars, 100μm (early) or 50μm (late). Data shown as mean ± SEM; each dot represents an individual mouse; analyzed by one way ANOVA followed by Tukey’s multiple comparison test. **(B**) Schematic of experimental paradigm for nasal anti-CD3 antibody administration to treat neuroinflammation in mild COVID mouse model. **(C)** Uniform manifold approximation and projection (UMAP) of scRNA-seq data in brain parenchyma cells enriched from subcortex region. **(D)** Distribution of different neuron subsets from the scRNAseq. **(E)** Upset plot comparing the microglia DEGs. **(F)** Volcano plot illustrating results of differential expression testing between microglia from iso compared with mock or anti-CD3. **(G-H)** GO pathways analysis comparing microglia DEGs between iso and mock or anti-CD3. **(I)** Quantified recognition ratio from novel objection recognition testing in 4 weeks antibody treated mild COVID or mock mice. **(J)** Schematic of the MY-LC study. **(K)** Percentage of circulating effector Treg cell and **(I)** nTreg in Long COVID patients with and without indicated symptoms. ns=no significant difference, \**p* <0.05, ** *p* <0.01, \*\*\**p* <0.001, \*\*\*\**p* <0.0001.

Because microglia can undergo substantial state transitions without changing in abundance (Fernandez-Castaneda et al., 2022), we next examined microglial transcriptional programs in the single cell dataset. Intersection analysis of microglial DEGs across contrasts showed that most transcriptional changes were specific to either infection (isotype vs mock) or treatment (aCD3 vs isotype), with a smaller shared gene set regulated in both comparisons (Figure 4E). Microglia from infected mice upregulated inflammatory and antigen presentation modules, whereas aCD3 induced a partially overlapping but distinct program (Figure 4F). Notably, the inhibitory receptor Cd300lf (Lu et al., 2024), which was elevated in microglia from isotype-treated mild COVID mice, was reduced by aCD3, with the largest shift occurring within the homeostatic microglia cluster as previously reported (Figure S2E, S2F)(Fernandez-Castaneda et al., 2022; Hammond et al., 2019), suggesting restoration of a more surveillance-like state. Pathway enrichment of microglial genes upregulated by aCD3 highlighted chemokine signaling/chemotaxis, phagosome and antigen-processing modules, and regulatory signaling pathways including TGF-β–related terms, consistent with enhanced immune regulation and debris clearance (Figure 4G). In contrast, genes downregulated by aCD3 were enriched for leukocyte activation, NF-κB, and cytokine-mediated inflammatory signaling pathways, indicating suppression of proinflammatory microglial programs (Figure 4H).

We then tested whether these cellular and molecular changes coincided with improved cognition. Using a modified novel object recognition test with a short time interval (5 min) to probe attention and short term memory relevant to patient reported “brain fog”(Gibson et al., 2019), isotype-treated mice at 8 weeks post infection failed to discriminate between familiar and novel objects (Fig. 4I). In contrast, mild COVID mice treated with nasal aCD3 for 4 weeks showed a restored preference for the novel object comparable to mock-infected animals, indicating rescue of attention and short-term recognition memory (Figure 4I).

Given the Treg-biased mechanism of nasal aCD3, we asked whether circulating Treg programs are linked to cognitive symptoms in people with Long COVID. Re-analysis of our MY-LC cohort (Klein et al., 2023), which includes symptom surveys and PBMC profiling (Figure 4J), showed that participants with Long COVID had neurological symptoms, including the cognitive complaints like brain fog, dizziness, and memory problems, consistently exhibited reduced circulating effTreg (Figure 4K) or nTreg (Figure 4L) compared with those without these symptoms. Together, these data indicate that nasal aCD3 ameliorates persistent deficits in hippocampal neurogenesis and short-term memory after mild respiratory SARS CoV-2 infection in mice, and that reduced effector Treg levels in circulation are associated with cognitive symptoms reported by patients with Long COVID, suggesting the role of Treg-mediated immune regulation in post-acute infection cognitive dysfunction.

## Discussion

Persistent cognitive dysfunction after SARS CoV-2 infection is a major symptom in Long COVID, and the biological circuits that remain amenable to intervention after symptoms are established are only beginning to be defined (Iwasaki and Putrino, 2023; Peluso and Deeks, 2024). Converging clinical and experimental data implicate neuroinflammation, including sustained microglial reactivity, as a key mediator of the neurological manifestations of Long COVID (Fernandez-Castaneda et al., 2022; Monje and Iwasaki, 2022). Here we show that nasal aCD3 mAb, delivered either early after infection or during an established post-acute phase, induces IL 10–producing Foxp3+ Treg cells, attenuates neuroinflammation, and restores hippocampal neurogenesis and cognitive behavior in a mild, respiratory-restricted COVID model.

Multiple lines of evidence support a role for sustained microglial reactivity in neurological Long COVID. Neuropathological and CSF studies have described microgliosis, astrogliosis, cytokine elevation, and blood–brain barrier disruption in patients with persistent neurological symptoms, including those who experienced only mild acute disease (Beiersdorfer et al., 2024; Farhadian et al., 2023; Grant et al., 2024; Greene et al., 2024). Mild COVID mouse models converge on this picture: mild, respiratory-restricted infection produces long-lasting microglial reactivity, myelin damage, and impaired hippocampal neurogenesis in the absence of direct viral neuroinvasion (Fernandez-Castaneda et al., 2022; Israelow et al., 2020). Microglia exhibited a consistent transcriptional signature weeks after infection, with upregulation of antigen presentation, interferon and NF-κB driven inflammatory pathways despite confinement of viral replication to the lung(Grant et al., 2024). Nasal aCD3 reshaped this landscape when given either early or after neuroinflammation was established, treatment reduced IBA1+ and GFAP+ cell densities in the hippocampus and white matter. Normalization of receptors such as Cd300lf within homeostatic microglia clusters (Fernandez-Castaneda et al., 2022; Keswani et al., 2020; Lu et al., 2024), together with suppression of inflammatory modules and enhancement of phagosome and TGF β–related pathways, is consistent with a shift from a chronically reactive, antigen-presenting state toward a surveillance-like phenotype more compatible with circuit homeostasis and repair.

Treg cells provide a mechanistic bridge between mucosal aCD3 mAb and these neuroimmune effects (Kuhn and Weiner, 2016). Tregs are central to restraining effector responses, enforcing tolerance at barrier sites and sculpting tissue resident macrophage and microglial states (Dikiy and Rudensky, 2023; Ito et al., 2019). Imbalanced Treg have been reported to be correlated with COVID-19 severity and Long COVID (Galvan-Pena et al., 2021; Shahbaz et al., 2025). In our model, nasal aCD3 expanded brain Foxp3+ CD4 T cells, including IL-10–producing and LAP expressing subsets associated with potent suppressive function, without compromising established antiviral humoral immunity as anti-spike antibody titer. This pattern argues that mucosal aCD3 does not broadly blunt antiviral immunity but instead skews T cell responses toward a regulatory phenotype capable of entering the CNS (Ito et al., 2019; Liston et al., 2024). The association between reduced circulating effector Tregs and cognitive symptom clusters in our Long COVID cohort provides a translational mirror, alongside additional inflammatory biomarkers such as CCL11 that have been linked to brain fog in prior studies (Fernandez-Castaneda et al., 2022; Geraghty et al., 2025), suggesting that patients with regulatory deficits may be particularly benefit from Treg-directed intervention.

At the neuronal level, nasal aCD3 rescued in hippocampal neurogenesis vulnerable to Long COVID. Mild respiratory infection reduced DCX+ neuroblasts in the dentate gyrus and impaired performance on a short delay novel object recognition task designed to probe aspects of “brain fog,” consistent with reports of long-lasting cognitive deficits and altered hippocampal structure and connectivity in human cohorts (Zorzo et al., 2023). aCD3 restored dentate gyrus neurogenesis across a broad temporal window and increased the representation of immature neuronal populations while reducing mature excitatory neurons, consistent with enhanced ongoing dentate neurogenesis and a shift in neuronal population balance (Disouky et al., 2026; Toni and Schinder, 2015). Prior work has implicated chemokines such as CCL11 as mediators of impaired neurogenesis and white matter microglial activation in different chronic CNS inflammation models (Fernandez-Castaneda et al., 2022; Geraghty et al., 2025; Villeda et al., 2011). In our delayed treatment paradigm, nasal aCD3 increased IL-10 and reduced CCL11 in the CSF and IL-10 production from Tregs, identifying a Treg–IL-10–CCL11 axis as one route by which immune rebalancing may release a brake on the neurogenic niche. Disentangling whether this axis acts directly on neural progenitors or indirectly via microglia, astrocytes or the neurovascular unit will require cell type specific manipulation of IL-10 and CCL11 signaling and longitudinal imaging of neurogenesis and myelin integrity.

The nasal route offers practical and biological advantages for inducing this regulatory circuit (Wu et al., 2008). The mucosal immune system is naturally biased toward tolerogenic responses, and nasal aCD3 has been shown to expand IL 10–producing Tregs that home to the CNS and modulate microglial activation in models of multiple sclerosis, Alzheimer’s disease and traumatic brain injury (Izzy et al., 2025; Lopes et al., 2023; Mayo et al., 2016). Foralumab, a fully human aCD3 formulated for nasal delivery, has been administered to people and produced immunomodulatory effects with a favorable safety profile (Kuhn and Weiner, 2016; Singhal et al., 2025), including a pilot trial in mild to moderate COVID 19 in which nasal foralumab reduced lung inflammation and circulating inflammatory biomarkers without major adverse events (Moreira et al., 2023; Moreira et al., 2021). Studies in progressive multiple sclerosis and have demonstrated nasal foralumab dampens microglia reactivity as measured by TSPO-PET imaging, modulates the immune system towards a regulatory phenotype and stabilizes disease(Chitnis et al., 2026; Tay et al., 2026), further supporting the feasibility of nasal aCD3 as a CNS targeted immunotherapy. Our findings here extend this concept to the post-acute phase of SARS CoV-2 infection, suggesting that nasal aCD3 could provide a noninvasive means to modulate brain relevant immune circuits in Long COVID.

Together, our findings identify nasal aCD3 as a candidate immunotherapy for Long COVID–associated cognitive dysfunction that operates by inducing IL-10–competent Tregs, reprogramming microglial states, normalizing chemokine milieus such as the CCL11 axis and restoring hippocampal neurogenesis and a cognitive behavioral indicator of attention and short-term memory. By integrating mechanistic mouse experiments with immune phenotyping in Long COVID patients, this work supports a broader framework in which a subtype of Long COVID with neuroimmune symptoms, characterized by effector Treg deficiency and persistent microglial activation, could be targeted with biomarker-anchored interventions in post-viral neuroinflammatory disease.

This study has several limitations. First, our mouse model was performed in female mice, a choice motivated by reports of higher Long COVID symptom burden in women(Silva et al., 2024), but it does not fully recapitulate the heterogeneous pathophysiology of human Long COVID. Second, our mechanistic investigations identified the Treg-IL-10-CCL11 axis as a key mediator of therapeutic benefit, but cell-type-specific loss-of-function studies will be required to definitively establish whether this axis acts directly on neural progenitors or indirectly via microglia, astrocytes, or the neurovascular unit. Third, cognitive assessment was limited to a single behavioral paradigm (the modified novel object recognition test), which evaluated short-term recognition memory and did not encompass other domains, such as executive function, and processing speed, because behavioral studies were constrained by the requirements of BSL3 facility. Finally, our human cohort analysis demonstrated associations between reduced circulating effector Tregs and cognitive symptoms in Long COVID patients, but these data are observational and do not establish causation. Despite these limitations, our findings provide a strong preclinical rationale to advance nasal aCD3 mAb therapy toward clinical evaluation for Long COVID-associated cognitive dysfunction.

## Methods

### Animal

6-to 12-week-old female CD1 mice were purchased from Charles River and housed at Yale University. Experiments with wild-type mice transduced with AAV-hACE2 were performed with littermate controls. All procedures used in this study (sex and age matched) complied with federal guidelines and the institutional policies of the Yale School of Medicine Animal Care and Use Committee.

### Virus Stock

As reported previously (Israelow et al., 2020), Vero E6 cells overexpressing angiotensin-converting enzyme 2 (ACE2) and TMPRSS2 [kindly provided by B. Graham at the National Institutes of Health Vaccine Research Center (NIH-VRC)] were infected with SARS-CoV-2 isolate hCOV-19/USA-WA1/2020 (NR-52281; BEI Resources) at a M.O.I. of 0.01 for 2 days until sufficient cytopathic effect was noted. After incubation, the supernatant was clarified by centrifugation (5 min, 500g), filtered through a 0.45-μm filter. Virus containing supernatant was applied to Amicon Ultra-15 centrifugal filter (Ultracel 100k) and spun at 2000 rpm for 15 minutes prior to aliquoting and storage at −80°C. Viral titers were measured with a standard plaque assay by using Vero E6 cells.

### AAV preparation

Adeno-associated virus 9 encoding hACE2 (AAV-CMV-hACE2) were made through the Penn vector core at University of Pennsylvania.

### AAV delivery and SARS-CoV2 infection

To overexpress human ACE2 in the lower respiratory tract, CD1 mice were infected intratracheally with AAV-hACE2 as previously described (Israelow et al., 2020). Briefly, animals were anaesthetized using a mixture of ketamine (100 mg/kg) and xylazine (10 mg/kg), injected intraperitoneally. The rostral neck was shaved and disinfected with iodine and ethanol. A 5-mm incision was made, the salivary glands were retracted, and the trachea was visualized. Using a 500-μl insulin syringe, a 50-μl bolus injection of 10^11^ genome copies (GC) of AAV-hACE2 was injected into the trachea. The incision was closed with VetBond skin glue. After two weeks of recovery, mice were anesthetized using 30% vol/vol isoflurane diluted in propylene glycol. Using a pipette, 50 µl of SARS-CoV-2 (3 × 10^7^ PFU/ml) was delivered intranasally

### Treatment with anti-CD3 mAb

Mice were nasally treated with a daily antibody dose, starting either one week or four weeks after SARS-CoV-2 infection for seven days. After one week of daily treatment, mice were treated with the antibody every other day for an additional three weeks. Each mouse was given 1 µg of hamster anti-mouse CD3 antibody (145-2C11, BioXCell) or hamster IgG control antibody (BioXCell)(Izzy et al., 2025).

### Flowcytometry

Brains were harvested from PBS perfusion mice and incubated in a digestion buffer containing 1 mg/ml collagenase D (Roche) and 16 μg/ml DNase I (Sigma-Aldrich) in HBSS at 37 °C for 30 min. Digested cell suspension was filtered through a 70-μm filter, followed by gradient centrifugation enrichment with 30% Percoll. Enrihched brain cells were blocked with Fc blocker (2.4G2, BioxCell) for 10 minutes on ice and then stained with aqua Live/Dead, Pacific blue anti-mouse CD45 (30-F11, BioLegend), APC/cyanine7 anti-mouse CD11b (M1/70, BioLegend), BV605 anti-mouse TCRβ (H57-597, BioLegend), APC anti-mouse CD4 (RM4-5, BioLegend) and PE anti-mouse LAP (TW7-16B4, BioLegend) for 30 minutes on ice. Cells were washed to remove excess antibodies and fixed with 4% PFA for 30 minutes before analysis. For intracellular staining, surface-stained cells were fixed and permeabilized with the FOXP3/Transcription Factor Staining Buffer Set (Thermo Fisher), followed by FITC anti-mouse Foxp3 staining (FJK-16s, Thermo Fisher). Cells were washed and resuspended in PBS for analysis on an Attune NXT (Thermo Fisher). Data were analyzed using FlowJo software version 10.8 software (BD)

### Cytokine/Chemokine screening

The Luminex multiplex assay was performed by the Human Immune Monitoring Center at Stanford University with Mouse 48-plex Procarta kits from Thermo Fisher as previously described(Fernandez-Castaneda et al., 2022). CSF samples were measured as singlets, while serum samples were measured in duplicate.

### Immunohistochemistry

Mice were deeply anesthetized with isoflurane until respiration rate slowed and perfused transcardially with Hank’s Balanced Salt Solution (HBSS). Brains were isolated and fixed in 4% paraformaldehyde (PFA) for 24-48 hours at 4°C. Following fixation, samples were sequentially cryoprotected by incubation in 30% sucrose solution for 24 h. Brains were then embedded in Tissue-Tek OCT compound (Sakura, compound 4583), snap-frozen, and preserved at −80 °C until further processing. Brains were subsequently sectioned at −20 °C using a cryostat at the bregma position for each targeted brain. Sections were cut in 0.2 mm in a sixfold series interval. Five total sections were placed on Colorfrost Plus-treated adhesion slides (Thermo Fisher Scientific) and stored at −20 °C until the time of staining. For immunofluorescent detection, sections were first allowed to equilibrate to room temperature, then permeabilized and blocked for 1 hour in a 10% normal horse serum solution, containing 0.1% Triton X-100, 1% glycine, and 2% bovine serum albumin (BSA). Slides were incubated overnight at 4 °C with anti-Iba1 (rabbit, 1:500, Wako), DCX (rabbit, 1:200, Abcam, cat.no. ab18723), or GFAP-AF488 conjugated antibody (mouse, 1:500, Thermo fisher, cat.no. 53-9892-82). For unconjugated primary antibodies, the following day the sections were washed and incubated with an Alexa Fluor-647 goat-anti rabbit IgG (1:1000, Thermo Fisher, cat.no. A-21245) or Alexa Fluor-594 goat anti-rabbit IgG (1:1000, Thermo Fisher, cat.no. A-11012). Stained slides were co-stained with DAPI mounting medium (Vector laboratories, cat. no. UX-93952-24). Images were taken using an Olympus FV3000 series confocal laser scanning microscope system on the 20x or 10x objective.

### Image analysis

Quantification of Iba-1+, GFAP+, and DCX+ cells was performed using ImageJ (https://imagej.nih.gov/ij). For each section, three representative photomicrographs were analyzed (3-4 sections per animal), and cell counts were averaged per field of view. Regions of interest (ROIs) were defined as 100 x 100 pixels, corresponding to a spatial resolution of 0.625 µm per pixel. Numbers are represented as cells per mm2.

### Behavioral testing

All behavioral assays were performed in a clean, empty rat cage (26.6 x 48.26 x 20.32 cm) covered with white plastic wrap and placed in an ABSL3 level Biosafety Cabinet. Animals were handled for 2 minutes for 4 days before subjecting to novel object recognition testing (NORT). One day before the NORT, the mouse was placed in the center of the rat cage and allowed to habituate for 20 min. The NORT is a modified version with a shortened rest window between the training and testing phase to evaluate the short-term memory (< 5 min) function. During the task, the mouse was placed in the center of the cage to acclimate for 10 min before being returned to the home cage for another 5 min. The mouse was then placed in the cage with two identical Lego objects for 5 min exploration and then returned to the home cage for 5 min. During this time, one object from the training phase was replaced a new Lego object (of similar size, different shape and color) for the testing phase. In the novel object testing phase, the mouse was returned to the cage and allowed to explore for 10 min. The recorded video was analyzed using Noldus analysis software (EthoVision XT 15). Only animals that explored both objects during the testing phase for an accumulation of a minimum of 20s were included in the analysis.

### Sample preparation of Bulk RNAseq

Mice were anesthetized with isoflurane and perfused intracardially with 10ml cold perfusion buffer. Perfusion buffer, dissection buffer and digestion buffer were freshly prepared as previously described(Marsh et al., 2022). Brains were quickly dissected and placed into an adult mouse brain slicer matrix within dissection buffer. 3μm cerebrum slice was cut and contained cortex and corpus callosum followed by chopping in digestion buffer. Chopped tissues were incubated for 45 minutes at 37°C and triturated by pipetting every 15 minutes. After incubation, the cell suspension was then passed through a 70μm filter. Cells were wash with completed DMEM and spun to pellet cells. Cell pellets were resuspended in 40% Percoll and followed by spinning for 25 minutes at 650g. The supernatant containing myelin debris was discarded and the pellet was resuspended in FACS buffer (HBSS+1% FBS+1mM EDTA) with inhibitory cocktail (Actinomycin D+Triptolide+Anisomycin).

For FACS sorting, the cells were resuspended in FC blocker solution (2.4G2, BioxCell) and incubated on ice for 10 mins. Cocktails of desired staining antibodies including live-dead Aqua, PE-anti-CD11b (M1/70, BioLegend) and Alexafluor647 anti-CD45 (30-F11, BioLegend) were added directly to cell suspension for 20 minutes incubation at 4°C. Prior to sort, cell were washed and resuspend with FACS buffer. Stained cells were sorted by BD FACSAria (BD Biosciences). RNA were extracted from sorted cells via the Qiagen Kit and submitted to Yale Center for Genome Analysis, who then created cDNA libraries and sequenced on IlluminaSeq at 100 million reads per sample.

All procedures except RNA extraction and sequencing were processed in Yale BSL-3 facility.

### Sample preparation and data analysis of scRNAseq

Single cell suspension were prepared as the samples for bulk RNAseq and loaded onto the Chromium Controller (10x Genomics) for droplet formation. scRNA-seq libraries were prepared using the Chromium Single Cell 3’Reagent Kit (10x Genomics), according to manufacturer’s protocol. Samples were sequenced on the NovaSeq 6000 with 28-bp read 1, 8-bp i7 index and 98-bp read 2 for the gene-expression library. Sequencing results were demultiplexed into Fastq files using the Cell Ranger (10x Genomics, 6.1.2) mkfastq function. Samples were aligned to mm10-2.2.0 10x genome. All procedures except sequencing were processed in Yale BSL-3 facility.

### Bioinformatics analysis

#### Mapping of bulk RNA-seq libraries and differential gene expression analysis

Initially raw reads served as input for Trimmomatic 0.40 (Bolger et al., 2014), which performed quality filtering removing Illumina adaptor sequences, low quality bases (phred score quality > 20), and short reads (PE -phred33 ILLUMINACLIP:truseq.fa:2:30:10 LEADING:3 TRAILING:3 SLIDINGWINDOW:3:20 MINLEN:36). Trimming was followed by rRNA filtering by SortMeRNA software (Kopylova et al., 2012) using the SILVA rRNA library (Yilmaz et al., 2014) as reference. Next the reads were mapped against the Genome Reference Consortium Mouse Build 39 (GRCm39) and the transcript abundance was quantified using Salmon v1.10.2 (Patro et al., 2017). Finally, the count data were direct to differential analysis with DESeq2 (version 1.48.1) R 4.5 package (Love et al., 2014). The Cook’s distance together with Principal components analysis and hierarchical clustering of the 1000 most variable genes were used to identify outliers.

#### Mouse brain scRNA-seq analysis

Reads were aligned to the mouse genome (refdata-gex-mm10-2020-A) using Cell Ranger software (v.7.0.1) (10x Genomics) generating the raw gene counts matrix. These count matrixes served as input to downstream analysis in Seurat (version 5.1.0) (Hao et al., 2021), initially filtering cells with unique molecular identifiers over 5,000 or less than 500, besides outliers with more than 5% mitochondrial counts. The feature expression measurements for each cell were normalized by the total expression, multiplied by the scale factor (10,000), and log transformed. We calculated the 3,000 highly variable genes to be used as integration anchors as we merged the three cohorts, followed by linear transformation (scaling) regressing out heterogeneity associated with mitochondrial contamination and UMI counts. Linear dimensional reduction was performed by PCA, followed by non-linear UMAP embedding using the first 30 principal components, same parameter used for clustering with FindNeighbors and FindClusters (0.9 resolution) Seurat’s functions. Differential gene-expression comparison between groups was done based on the non-parametric Wilcoxon rank sum test. Genes with fold change > 1.5 (absolute value), adjusted P value (Bonferroni correction) < 0.1 and detected in a minimum fraction of 20% cells in either of the two populations were considered differentially expressed.

#### Pathway analysis

Biological process and Gene Ontology terms search and analysis were performed through online database PANTHER 19.0 (Mi et al., 2019), Enrichr-KG (Chen et al., 2013; Kuleshov et al., 2016; Xie et al., 2021), Metascape (Zhou et al., 2019) and WebGestalt (Elizarraras et al., 2024) with standard parameters.

## Data and code availability

The bulk RNAseq and scRNAseq raw data presented in this work will be publicly available at the Sequence Read Archive (SRA) after publication. Bulk and scRNA-seq data is submitted to SRA. with the following accession codes.

## Acknowledgements

The authors would like to thank members of the Yale Center for Genomic Analysis (YCGA) and the Rodent Behavior Analysis facility at Yale Kavli Institute for Neuroscience. This study was funded by National Institutes of Health (K08NS123503 to S.I.), Stepping Strong Breakthrough Award from Gillian Reny Stepping Strong Center for Trauma Innovation (to S.I.), Department of Defense (HT9425-24-1-0635, W911NF-23-1-0276, and SC240188 to S.I.), the Howard Hughes Medical Institute Emerging Pathogens Initiative (to M.M. and A.I.), the Howard Hughes Medical Institute (to M.M. and A.I.), and the Else Kröner Fresenius Prize for Medical Research 2023 (to A.I.).

## Author Contributions

P.L., S.I., A.I. and H.L.W. planned the project; P.L. and S.I. designed, analyzed and interpreted data and wrote the manuscript; P.L., S.I., T.Y., H.T.I., J.R.C., P.D.S., M.H.A.M., A.A. performed experiments and analyzed data; review and editing by P.L., S.I., T.Y., H.T.I., J.R.C., P.D.S., M.H.A.M., A.A., T.G.M., M.M., A.I. and H.L.W.; supervised by S.I., A.I. and H.L.W.; funding acquisition by S.I., M. M, A.I. and H.L.W..

## Declaration of Conflict of Interest

H.L.W. is on the scientific advisory board of Tiziana Life Sciences. A.I. co-founded RIGImmune, Rho Bio., Xanadu Bio, PanV, and is a member of the Board of Directors of Roche Holding Ltd and Genentech. All other authors declare no conflict of interest.

---

**Figure. S1.**
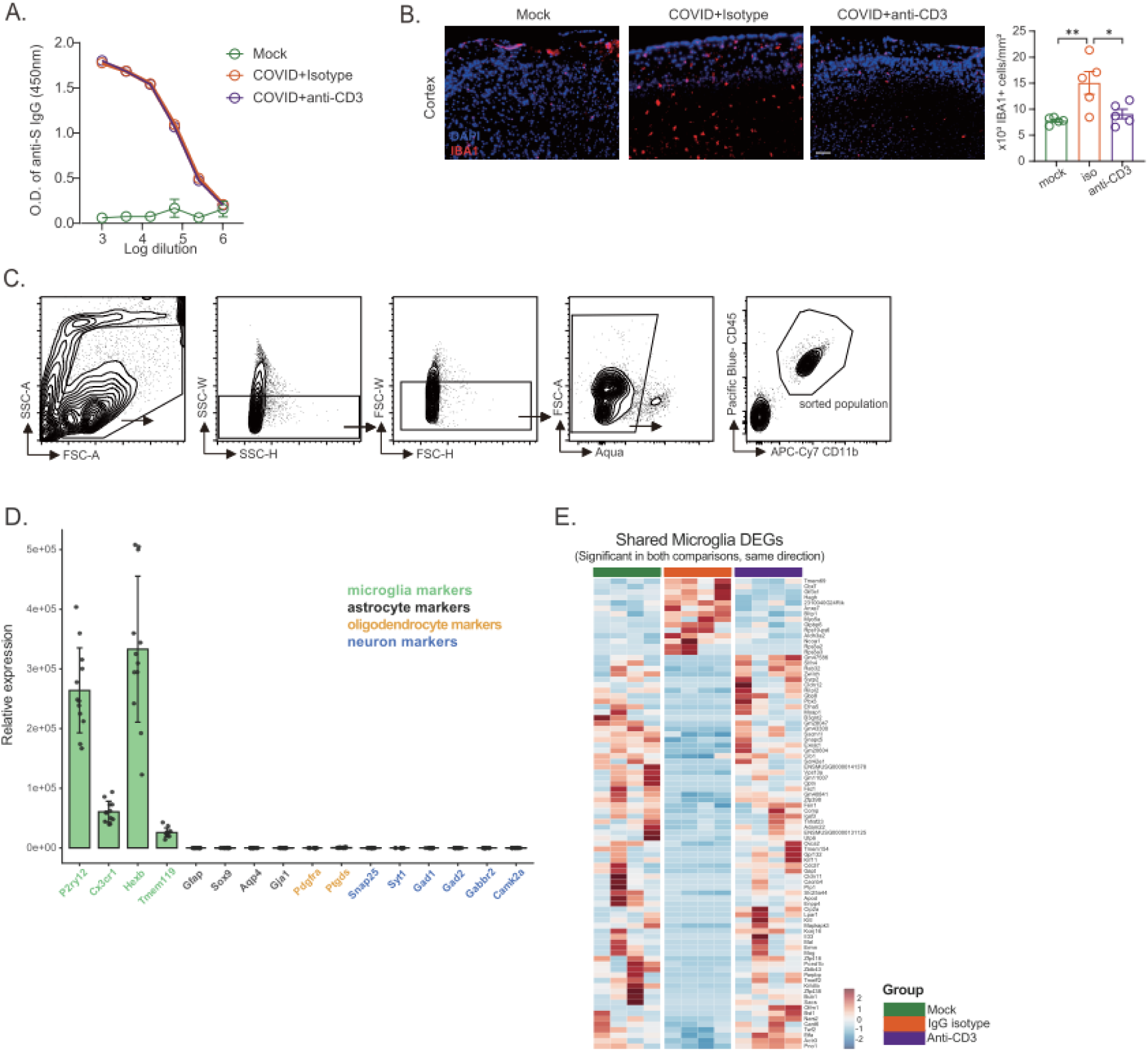
Additional data of early nasal anti-CD3 treatment in mild COVID mice, related to Figure1 & 2. **(A**) Serum antibodies were measured against spike protein using an ELISA, n=6-7 mice per group. (**B)** Quantification and confocal micrograph of microglia (IBA1+) in the cortex after 4 weeks isotype or anti-CD3 antibody administration in mild COVID or mock mice; n=6-7 mice per group. Data shown as mean ± SEM; each dot represents an individual mouse. *p <0.05, ** p <0.01, ***p <0.001, ****p <0.0001 (one way ANOVA followed by Tukey’s multiple comparison test). **(C)** Representative gating strategy to isolate microglia for bulk RNAseq. **(D)** Relative expression of cell type-specific markers in sorted microglia. **(E)** Heatmap of shared microglial DEGs significant in both comparisons (Mock vs IgG isotype and anti-CD3 vs IgG isotype) and changing in the same direction. n=4 or 5 mice per group.

**Figure. S2.**
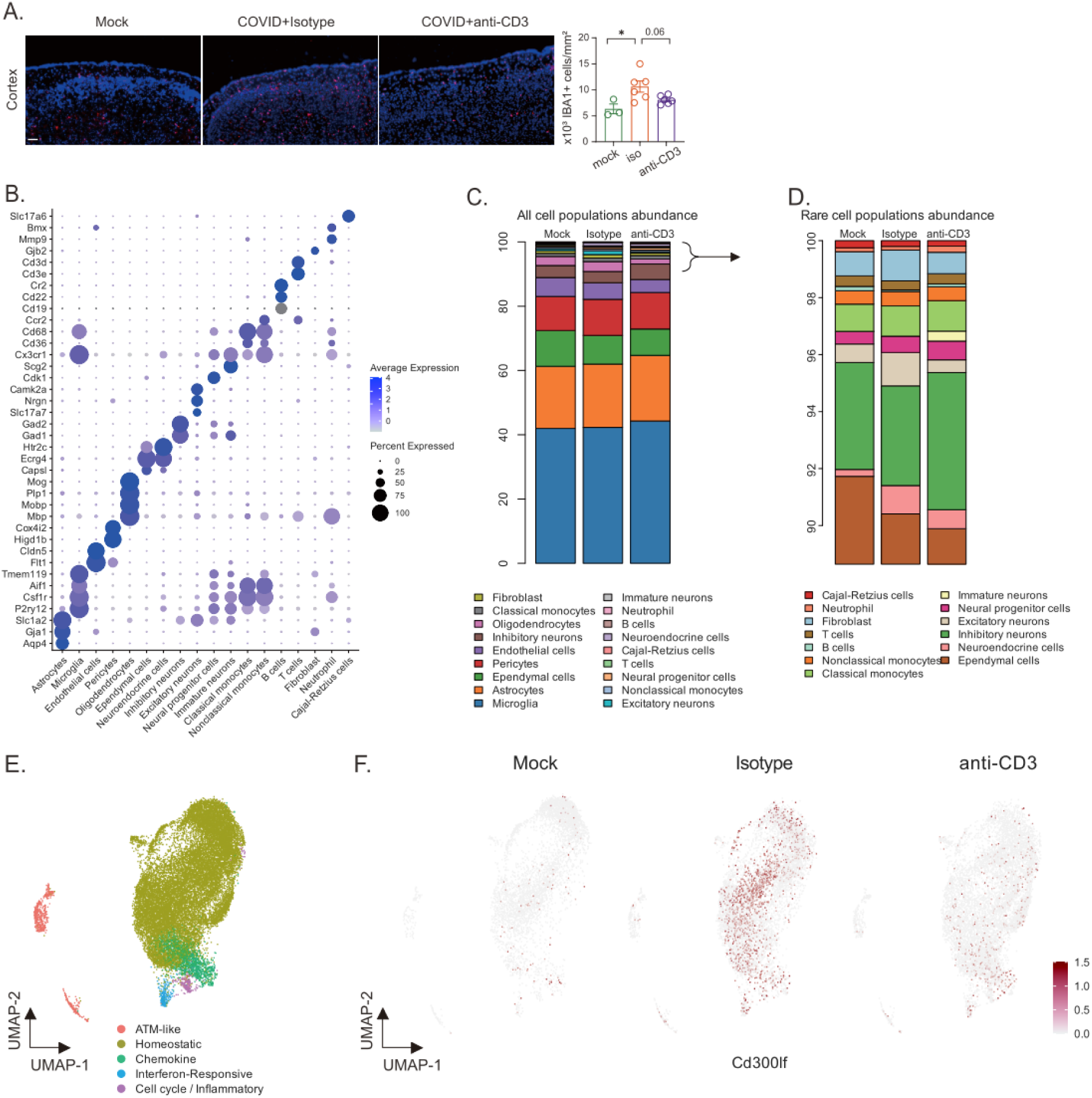
Additional data of late nasal anti-CD3 treatment in mild COVID mice, related to Figure3 & 4. **(A**) Quantification and confocal micrograph of microglia (IBA1+, red) in the cortex after 4 weeks isotype or anti-CD3 antibody administration in mild COVID or mock mice; n=3-7 mice per group. Data shown as mean ± SEM; each dot represents an individual mouse. *p <0.05, ** p <0.01, ***p <0.001, ****p <0.0001 (one way ANOVA followed by Tukey’s multiple comparison test) .(**B)** Dot plot showing expression of cell type marker genes used in the annotation the clusters shown in figure 4C and 4D. **(C)** Distribution of total cell subsets in the scRNAseq and **(D)** distribution of the least abundant cell types. **(E)** UMAP plot of microglia colored by clusters. **(F)** UMAP plots as in (E) colored by expression level of Cd300lf.

## Notes

### Summary of Updates

No content changed. Copyright updated to CC BY 4.0 for maximal accessibility.

## References

Al-Aly, Z., Davis, H., McCorkell, L., Soares, L., Wulf-Hanson, S., Iwasaki, A., and Topol, E.J. (2024). Long COVID science, research and policy. Nat Med 30, 2148–2164.

Beiersdorfer, A., Rotermund, N., Schulz, K., Busch, M., Hirnet, D., Henne, S., Schwarzenberg, F.L., Dottermusch, M., Ondruschka, B., Matschke, J., et al. (2024). Microgliosis, astrogliosis and loss of aquaporin-4 polarity in frontal cortex of COVID-19 patients. bioRxiv.

Bolger, A.M., Lohse, M., and Usadel, B. (2014). Trimmomatic: a flexible trimmer for Illumina sequence data. Bioinformatics 30, 2114–2120.

Bonhenry, D., Charnley, M., Goncalves, J., Hammarstrom, P., Heneka, M.T., Itzhaki, R., Lambert, J.C., Mannan, M., Baig, A.M., Middeldorp, J., et al. (2024). SARS-CoV-2 infection as a cause of neurodegeneration. Lancet Neurol 23, 562–563.

Bonilla, H., Peluso, M.J., Rodgers, K., Aberg, J.A., Patterson, T.F., Tamburro, R., Baizer, L., Goldman, J.D., Rouphael, N., Deitchman, A., et al. (2023). Therapeutic trials for long COVID-19: A call to action from the interventions taskforce of the RECOVER initiative. Front Immunol 14, 1129459.

Braga, J., Lepra, M., Kish, S.J., Rusjan, P.M., Nasser, Z., Verhoeff, N., Vasdev, N., Bagby, M., Boileau, I., Husain, M.I., et al. (2023). Neuroinflammation After COVID-19 With Persistent Depressive and Cognitive Symptoms. JAMA Psychiatry 80, 787–795.

Bruno, R.M., Badhwar, S., Abid, L., Agharazii, M., Anastasio, F., Bellien, J., Burghuber, O., Faconti, L., Filipovsky, J., Ghiadoni, L., et al. (2025). Accelerated vascular ageing after COVID-19 infection: the CARTESIAN study. Eur Heart J 46, 3905–3918.

Chatenoud, L., and Bluestone, J.A. (2007). CD3-specific antibodies: a portal to the treatment of autoimmunity. Nat Rev Immunol 7, 622–632.

Chen, E.Y., Tan, C.M., Kou, Y., Duan, Q., Wang, Z., Meirelles, G.V., Clark, N.R., and Ma’ayan, A. (2013). Enrichr: interactive and collaborative HTML5 gene list enrichment analysis tool. BMC Bioinformatics 14, 128.

Chitnis, T., Singhal, T., Zurawski, J., Saraceno, T.J., Gopalakrishnan, N., Cain, L., LaBarre, B., King, D., Bergmark, R.W., Maxfield, A.Z., et al. (2026). Nasal Foralumab for the Treatment of Progression Independent of Relapses in Patients With Nonactive Secondary Progressive Multiple Sclerosis. Neurology Neuroimmunology & Neuroinflammation 13, e200543.

Davis, H.E., McCorkell, L., Vogel, J.M., and Topol, E.J. (2023). Long COVID: major findings, mechanisms and recommendations. Nat Rev Microbiol 21, 133–146.

Dikiy, S., and Rudensky, A.Y. (2023). Principles of regulatory T cell function. Immunity 56, 240–255.

Disouky, A., Sanborn, M.A., Sabitha, K.R., Mostafa, M.M., Ayala, I.A., Bennett, D.A., Lu, Y., Zhou, Y., Keene, C.D., Weintraub, S., et al. (2026). Human hippocampal neurogenesis in adulthood, ageing and Alzheimer’s disease. Nature.

Elizarraras, J.M., Liao, Y., Shi, Z., Zhu, Q., Pico, A.R., and Zhang, B. (2024). WebGestalt 2024: faster gene set analysis and new support for metabolomics and multi-omics. Nucleic Acids Res 52, W415–W421.

Ellen, A. (2025). From Stagnation to Strategy: Challenges in Advancing Long COVID Research. J Eval Clin Pract 31, e70180.

Eng, L.F., Vanderhaeghen, J.J., Bignami, A., and Gerstl, B. (1971). An acidic protein isolated from fibrous astrocytes. Brain Res 28, 351–354.

Farhadian, S.F., Reisert, H.D., McAlpine, L., Chiarella, J., Kosana, P., Yoon, J., and Spudich, S. (2023). Self-Reported Neuropsychiatric Post-COVID-19 Condition and CSF Markers of Neuroinflammation. JAMA Netw Open 6, e2342741.

Fernandez-Castaneda, A., Lu, P., Geraghty, A.C., Song, E., Lee, M.H., Wood, J., O’Dea, M.R., Dutton, S., Shamardani, K., Nwangwu, K., et al. (2022). Mild respiratory COVID can cause multi-lineage neural cell and myelin dysregulation. Cell 185, 2452–2468 e2416.

Fujimoto, Y., Abe, H., Eiro, T., Tsugawa, S., Tanaka, M., Hatano, M., Nakajima, W., Ichijo, S., Arisawa, T., Takada, Y., et al. (2025). Systemic increase of AMPA receptors associated with cognitive impairment of long COVID. Brain Commun 7, fcaf337.

Galvan-Pena, S., Leon, J., Chowdhary, K., Michelson, D.A., Vijaykumar, B., Yang, L., Magnuson, A.M., Chen, F., Manickas-Hill, Z., Piechocka-Trocha, A., et al. (2021). Profound Treg perturbations correlate with COVID-19 severity. Proc Natl Acad Sci U S A 118.

Gandhi, R., Farez, M.F., Wang, Y., Kozoriz, D., Quintana, F.J., and Weiner, H.L. (2010). Cutting edge: human latency-associated peptide+ T cells: a novel regulatory T cell subset. J Immunol 184, 4620–4624.

Geraghty, A.C., Acosta-Alvarez, L., Rotiroti, M.C., Dutton, S., O’Dea, M.R., Kim, W., Trivedi, V., Mancusi, R., Shamardani, K., Malacon, K., et al. (2025). Immunotherapy-related cognitive impairment after CAR T cell therapy in mice. Cell 188, 3238–3258 e3225.

Gibson, E.M., Nagaraja, S., Ocampo, A., Tam, L.T., Wood, L.S., Pallegar, P.N., Greene, J.J., Geraghty, A.C., Goldstein, A.K., Ni, L., et al. (2019). Methotrexate Chemotherapy Induces Persistent Tri-glial Dysregulation that Underlies Chemotherapy-Related Cognitive Impairment. Cell 176, 43–55 e13.

Grant, R.A., Poor, T.A., Sichizya, L., Diaz, E., Bailey, J.I., Soni, S., Senkow, K.J., Perez-Leonor, X.G., Abdala-Valencia, H., Lu, Z., et al. (2024). Prolonged exposure to lung-derived cytokines is associated with activation of microglia in patients with COVID-19. JCI Insight *9*.

Green, R., Marjenberg, Z., Lip, G.Y.H., Banerjee, A., Wisnivesky, J., Delaney, B.C., Peluso, M.J., Wynberg, E., and Abduljawad, S. (2025). A systematic review and meta-analysis of the impact of vaccination on prevention of long COVID. Nat Commun 16, 10326.

Greene, C., Connolly, R., Brennan, D., Laffan, A., O’Keeffe, E., Zaporojan, L., O’Callaghan, J., Thomson, B., Connolly, E., Argue, R., et al. (2024). Blood-brain barrier disruption and sustained systemic inflammation in individuals with long COVID-associated cognitive impairment. Nat Neurosci 27, 421–432.

Groff, D., Sun, A., Ssentongo, A.E., Ba, D.M., Parsons, N., Poudel, G.R., Lekoubou, A., Oh, J.S., Ericson, J.E., Ssentongo, P., et al. (2021). Short-term and Long-term Rates of Postacute Sequelae of SARS-CoV-2 Infection: A Systematic Review. JAMA Netw Open 4, e2128568.

Hamlin, R.E., and Blish, C.A. (2024). Challenges and opportunities in long COVID research. Immunity 57, 1195–1214.

Hammond, T.R., Dufort, C., Dissing-Olesen, L., Giera, S., Young, A., Wysoker, A., Walker, A.J., Gergits, F., Segel, M., Nemesh, J., et al. (2019). Single-Cell RNA Sequencing of Microglia throughout the Mouse Lifespan and in the Injured Brain Reveals Complex Cell-State Changes. Immunity 50, 253–271 e256.

Hampshire, A., Trender, W., Chamberlain, S.R., Jolly, A.E., Grant, J.E., Patrick, F., Mazibuko, N., Williams, S.C., Barnby, J.M., Hellyer, P., et al. (2021). Cognitive deficits in people who have recovered from COVID-19. EClinicalMedicine 39, 101044.

Hao, Y., Hao, S., Andersen-Nissen, E., Mauck, W.M., 3rd, Zheng, S., Butler, A., Lee, M.J., Wilk, A.J., Darby, C., Zager, M., et al. (2021). Integrated analysis of multimodal single-cell data. Cell 184, 3573–3587 e3529.

Hol, E.M., and Pekny, M. (2015). Glial fibrillary acidic protein (GFAP) and the astrocyte intermediate filament system in diseases of the central nervous system. Curr Opin Cell Biol 32, 121–130.

Israelow, B., Song, E., Mao, T., Lu, P., Meir, A., Liu, F., Alfajaro, M.M., Wei, J., Dong, H., Homer, R.J., et al. (2020). Mouse model of SARS-CoV-2 reveals inflammatory role of type I interferon signaling. J Exp Med 217.

Ito, M., Komai, K., Mise-Omata, S., Iizuka-Koga, M., Noguchi, Y., Kondo, T., Sakai, R., Matsuo, K., Nakayama, T., Yoshie, O., et al. (2019). Brain regulatory T cells suppress astrogliosis and potentiate neurological recovery. Nature 565, 246–250.

Iwasaki, A., and Putrino, D. (2023). Why we need a deeper understanding of the pathophysiology of long COVID. Lancet Infect Dis 23, 393–395.

Izzy, S., Yahya, T., Albastaki, O., Abou-El-Hassan, H., Aronchik, M., Cao, T., De Oliveira, M.G., Lu, K.J., Moreira, T.G., da Silva, P., et al. (2025). Nasal anti-CD3 monoclonal antibody ameliorates traumatic brain injury, enhances microglial phagocytosis and reduces neuroinflammation via IL-10-dependent T(reg)-microglia crosstalk. Nat Neurosci 28, 499–516.

Jansen, E.B., Orvold, S.N., Swan, C.L., Yourkowski, A., Thivierge, B.M., Francis, M.E., Ge, A., Rioux, M., Darbellay, J., Howland, J.G., et al. (2022). After the virus has cleared-Can preclinical models be employed for Long COVID research? PLoS Pathog 18, e1010741.

Keswani, T., Roland, J., Herbert, F., Delcroix-Genete, D., Bauderlique-Le Roy, H., Gaayeb, L., Cazenave, P.A., and Pied, S. (2020). Expression of CD300lf by microglia contributes to resistance to cerebral malaria by impeding the neuroinflammation. Genes Immun 21, 45–62.

Klein, J., Wood, J., Jaycox, J.R., Dhodapkar, R.M., Lu, P., Gehlhausen, J.R., Tabachnikova, A., Greene, K., Tabacof, L., Malik, A.A., et al. (2023). Distinguishing features of long COVID identified through immune profiling. Nature 623, 139–148.

Kopylova, E., Noe, L., and Touzet, H. (2012). SortMeRNA: fast and accurate filtering of ribosomal RNAs in metatranscriptomic data. Bioinformatics 28, 3211–3217.

Kuhn, C., and Weiner, H.L. (2016). Therapeutic anti-CD3 monoclonal antibodies: from bench to bedside. Immunotherapy 8, 889–906.

Kuleshov, M.V., Jones, M.R., Rouillard, A.D., Fernandez, N.F., Duan, Q., Wang, Z., Koplev, S., Jenkins, S.L., Jagodnik, K.M., Lachmann, A., et al. (2016). Enrichr: a comprehensive gene set enrichment analysis web server 2016 update. Nucleic Acids Res 44, W90–97.

Liston, A., Dooley, J., and Yshii, L. (2022). Brain-resident regulatory T cells and their role in health and disease. Immunol Lett 248, 26–30.

Liston, A., Pasciuto, E., Fitzgerald, D.C., and Yshii, L. (2024). Brain regulatory T cells. Nat Rev Immunol 24, 326–337.

Lopes, J.R., Zhang, X., Mayrink, J., Tatematsu, B.K., Guo, L., LeServe, D.S., Abou-El-Hassan, H., Rong, F., Dalton, M.J., Oliveira, M.G., et al. (2023). Nasal administration of anti-CD3 monoclonal antibody ameliorates disease in a mouse model of Alzheimer’s disease. Proc Natl Acad Sci U S A 120, e2309221120.

Love, M.I., Huber, W., and Anders, S. (2014). Moderated estimation of fold change and dispersion for RNA-seq data with DESeq2. Genome Biol 15, 550.

Lu, Z., Liu, Z., Wang, C., Jiang, R., Wang, Z., Liao, W., Wang, W., Chen, J., Zhu, X., Zhao, J., et al. (2024). CD300LF(+) microglia impede the neuroinflammation following traumatic brain injury by inhibiting STING pathway. CNS Neurosci Ther 30, e14824.

Marsh, S.E., Walker, A.J., Kamath, T., Dissing-Olesen, L., Hammond, T.R., de Soysa, T.Y., Young, A.M.H., Murphy, S., Abdulraouf, A., Nadaf, N., et al. (2022). Dissection of artifactual and confounding glial signatures by single-cell sequencing of mouse and human brain. Nat Neurosci 25, 306–316.

Mayo, L., Cunha, A.P., Madi, A., Beynon, V., Yang, Z., Alvarez, J.I., Prat, A., Sobel, R.A., Kobzik, L., Lassmann, H., et al. (2016). IL-10-dependent Tr1 cells attenuate astrocyte activation and ameliorate chronic central nervous system inflammation. Brain 139, 1939–1957.

Mi, H., Muruganujan, A., Huang, X., Ebert, D., Mills, C., Guo, X., and Thomas, P.D. (2019). Protocol Update for large-scale genome and gene function analysis with the PANTHER classification system (v.14.0). Nat Protoc 14, 703–721.

Monje, M., and Iwasaki, A. (2022). The neurobiology of long COVID. Neuron 110, 3484–3496.

Moreira, T., Gauthier, C.D., Murphy, L., Lanser, T.B., Paul, A., Matos, K.T.F., Mangani, D., Izzy, S., Rezende, R.M., Healy, B.C., et al. (2023). Nasal administration of anti-CD3 mAb (Foralumab) downregulates NKG7 and increases TGFB1 and GIMAP7 expression in T cells in subjects with COVID-19. Proc Natl Acad Sci U S A 120, e2220272120.

Moreira, T.G., Matos, K.T.F., De Paula, G.S., Santana, T.M.M., Da Mata, R.G., Pansera, F.C., Cortina, A.S., Spinola, M.G., Baecher-Allan, C.M., Keppeke, G.D., et al. (2021). Nasal Administration of Anti-CD3 Monoclonal Antibody (Foralumab) Reduces Lung Inflammation and Blood Inflammatory Biomarkers in Mild to Moderate COVID-19 Patients: A Pilot Study. Front Immunol 12, 709861.

Mukherjee, S., Singer, T., Venkatesh, A., Choudhury, N.A., Perez Giraldo, G.S., Jimenez, M., Miller, J., Lopez, M., Hanson, B.A., Bawa, A.P., et al. (2025). Vaccination prior to SARS-CoV-2 infection does not affect the neurologic manifestations of long COVID. Brain Commun 7, fcae448.

Nalbandian, A., Sehgal, K., Gupta, A., Madhavan, M.V., McGroder, C., Stevens, J.S., Cook, J.R., Nordvig, A.S., Shalev, D., Sehrawat, T.S., et al. (2021). Post-acute COVID-19 syndrome. Nat Med 27, 601–615.

Notarte, K.I., Catahay, J.A., Velasco, J.V., Pastrana, A., Ver, A.T., Pangilinan, F.C., Peligro, P.J., Casimiro, M., Guerrero, J.J., Gellaco, M.M.L., et al. (2022). Impact of COVID-19 vaccination on the risk of developing long-COVID and on existing long-COVID symptoms: A systematic review. EClinicalMedicine 53, 101624.

Ochi, H., Abraham, M., Ishikawa, H., Frenkel, D., Yang, K., Basso, A.S., Wu, H., Chen, M.L., Gandhi, R., Miller, A., et al. (2006). Oral CD3-specific antibody suppresses autoimmune encephalomyelitis by inducing CD4+ CD25-LAP+ T cells. Nat Med 12, 627–635.

Pagliano, P., Salzano, F., D’Amore, C., Spera, A., Conti, V., Folliero, V., Franci, G., and Ascione, T. (2025). How do drug discovery scientists address the unmet need of long COVID syndrome therapeutics and what more can be done? Expert Opin Drug Discov 20, 1251–1265.

Patro, R., Duggal, G., Love, M.I., Irizarry, R.A., and Kingsford, C. (2017). Salmon provides fast and bias-aware quantification of transcript expression. Nat Methods 14, 417–419.

Peluso, M.J., and Deeks, S.G. (2024). Mechanisms of long COVID and the path toward therapeutics. Cell 187, 5500–5529.

Rubtsov, Y.P., Rasmussen, J.P., Chi, E.Y., Fontenot, J., Castelli, L., Ye, X., Treuting, P., Siewe, L., Roers, A., Henderson, W.R., Jr., et al. (2008). Regulatory T cell-derived interleukin-10 limits inflammation at environmental interfaces. Immunity 28, 546–558.

Saikarthik, J., Saraswathi, I., Padhi, B.K., Shamim, M.A., Alzerwi, N., Alarifi, A., and Gandhi, A.P. (2025). Structural and functional neuroimaging of hippocampus to study adult neurogenesis in long COVID-19 patients with neuropsychiatric symptoms: a scoping review. PeerJ 13, e19575.

Seo, D., Choi, Y., Jeong, E., Bang, S., Lee, J.S., Jang, I.H., Choi, L., Kim, J.H., Shin, W., Seo, B.R., et al. (2025). Distinct brain alterations and neurodegenerative processes in cognitive impairment associated with post-acute sequelae of COVID-19. Nat Commun 16, 10552.

Shahbaz, S., Osman, M., Syed, H., Mason, A., Rosychuk, R.J., Cohen Tervaert, J.W., and Elahi, S. (2025). Integrated immune, hormonal, and transcriptomic profiling reveals sex-specific dysregulation in long COVID patients with ME/CFS. Cell Rep Med 6, 102449.

Silva, J., Takahashi, T., Wood, J., Lu, P., Tabachnikova, A., Gehlhausen, J.R., Greene, K., Bhattacharjee, B., Monteiro, V.S., Lucas, C., et al. (2024). Sex differences in symptomatology and immune profiles of Long COVID. medRxiv.

Singhal, T., Cicero, S., Gale, S.A., Horan, N., Dubey, S., Marshall, G.A., and Weiner, H.L. (2025). Dampening of Microglial Activation With Nasal Foralumab Administration in Moderate Alzheimer’s Disease Dementia. Clin Nucl Med 50, 756–757.

Tay, D., Ahmed, H., Dawoud, A., Salam, M., Gobbi, L., Grether, U., Edelmann, M.R., Wittwer, M.B., Collin, L., Atz, K., et al. (2026). Translational molecular imaging and drug development in multiple sclerosis. Theranostics 16, 1630–1657.

Toni, N., and Schinder, A.F. (2015). Maturation and Functional Integration of New Granule Cells into the Adult Hippocampus. Cold Spring Harb Perspect Biol 8, a018903.

Vanderheiden, A., Hill, J.D., Jiang, X., Deppen, B., Bamunuarachchi, G., Soudani, N., Joshi, A., Cain, M.D., Boon, A.C.M., and Klein, R.S. (2024). Vaccination reduces central nervous system IL-1beta and memory deficits after COVID-19 in mice. Nat Immunol 25, 1158–1171.

Vignali, D.A., Collison, L.W., and Workman, C.J. (2008). How regulatory T cells work. Nat Rev Immunol 8, 523–532.

Villeda, S.A., Luo, J., Mosher, K.I., Zou, B., Britschgi, M., Bieri, G., Stan, T.M., Fainberg, N., Ding, Z., Eggel, A., et al. (2011). The ageing systemic milieu negatively regulates neurogenesis and cognitive function. Nature 477, 90–94.

Weiner, H.L., da Cunha, A.P., Quintana, F., and Wu, H. (2011). Oral tolerance. Immunol Rev 241, 241–259.

Wu, H.Y., Quintana, F.J., and Weiner, H.L. (2008). Nasal anti-CD3 antibody ameliorates lupus by inducing an IL-10-secreting CD4+ CD25-LAP+ regulatory T cell and is associated with down-regulation of IL-17+ CD4+ ICOS+ CXCR5+ follicular helper T cells. J Immunol 181, 6038–6050.

Xie, Z., Bailey, A., Kuleshov, M.V., Clarke, D.J.B., Evangelista, J.E., Jenkins, S.L., Lachmann, A., Wojciechowicz, M.L., Kropiwnicki, E., Jagodnik, K.M., et al. (2021). Gene Set Knowledge Discovery with Enrichr. Curr Protoc 1, e90.

Xu, E., Xie, Y., and Al-Aly, Z. (2022). Long-term neurologic outcomes of COVID-19. Nat Med 28, 2406–2415.

Yilmaz, P., Parfrey, L.W., Yarza, P., Gerken, J., Pruesse, E., Quast, C., Schweer, T., Peplies, J., Ludwig, W., and Glockner, F.O. (2014). The SILVA and “All-species Living Tree Project (LTP)” taxonomic frameworks. Nucleic Acids Res 42, D643–648.

Yin, K., Peluso, M.J., Luo, X., Thomas, R., Shin, M.G., Neidleman, J., Andrew, A., Young, K.C., Ma, T., Hoh, R., et al. (2024). Long COVID manifests with T cell dysregulation, inflammation and an uncoordinated adaptive immune response to SARS-CoV-2. Nat Immunol 25, 218–225.

Zhou, Y., Zhou, B., Pache, L., Chang, M., Khodabakhshi, A.H., Tanaseichuk, O., Benner, C., and Chanda, S.K. (2019). Metascape provides a biologist-oriented resource for the analysis of systems-level datasets. Nat Commun 10, 1523.

Zorzo, C., Solares, L., Mendez, M., and Mendez-Lopez, M. (2023). Hippocampal alterations after SARS-CoV-2 infection: A systematic review. Behav Brain Res 455, 114662.

